# A view of bird’s eyes – Pigeons lock their eyes in place during flight

**DOI:** 10.64898/2026.01.07.698303

**Authors:** Ivo G. Ros, Partha S. Bhagavatula, Andrew A. Biewener

## Abstract

Vision in most animals follows a fixate-and-saccade pattern. Birds fixate their viewing direction, then rapidly shift this gaze through head and eye movements. We used a head-mounted eye-tracking system in flying pigeons to relate eye to head movement and map eye position within the head. After take-off, the birds increased pupil size and adopted a fixed and consistent eye position in their head. In different visual environments, eye position returned to within 1° during flight. When flying, the birds positioned their eyes close to the primary horizontal axes of their vestibular systems. Because visual neurons share a common reference frame with the vestibular system, a consistent flight gaze position may actively align vision with mechanosensation and facilitate perception of self-motion.

**One-Sentence Summary:** A head-mounted eye-tracking system shows that pigeons adopt a consistent eye-in-head position during free flight

## Introduction

Most animals view the world by looking in one direction before rapidly moving their eyes to another direction. They fixate their eyes to stabilize the retinal image and prevent motion blur (Yarbus, 1967). Alternating with these fixations, animals move their eyes rapidly, or saccade, to change viewing direction and limit periods of motion blur (Yarbus, 1967; Land, 1973, 1999).

This ancient fixate-and-saccade pattern evolved in fishes and, independently, in invertebrates such as crustaceans, insects, and cephalopods (Land, 2019; Walls, 1962). Together with primates, these animals tend to change their viewing direction, or gaze, by moving their eyes.

Rather than moving their eyes, birds predominantly change their gaze by moving their heads. Specifically, head rotations account for more than 80% of a bird’s gaze changes in standing or restrained birds presented with wide-field visual stimuli (Gioanni, 1988; Maurice and Gioanni, 2004; Moinard, 2005). Large eye movements, however, are primarily used for near-field tasks such as food pecking (Martinoya *et al*., 1984a). During flight, however, avian eye position remains uncharacterized. To address gaze control in birds, we developed a head-mounted system (Fig. 1) in free-flying pigeons that relates eye to head movement and maps eye position within the head. We hypothesize that free-flying birds hold their eyes fixed within their head, such that head movement controls gaze direction.

**Fig. 1.**
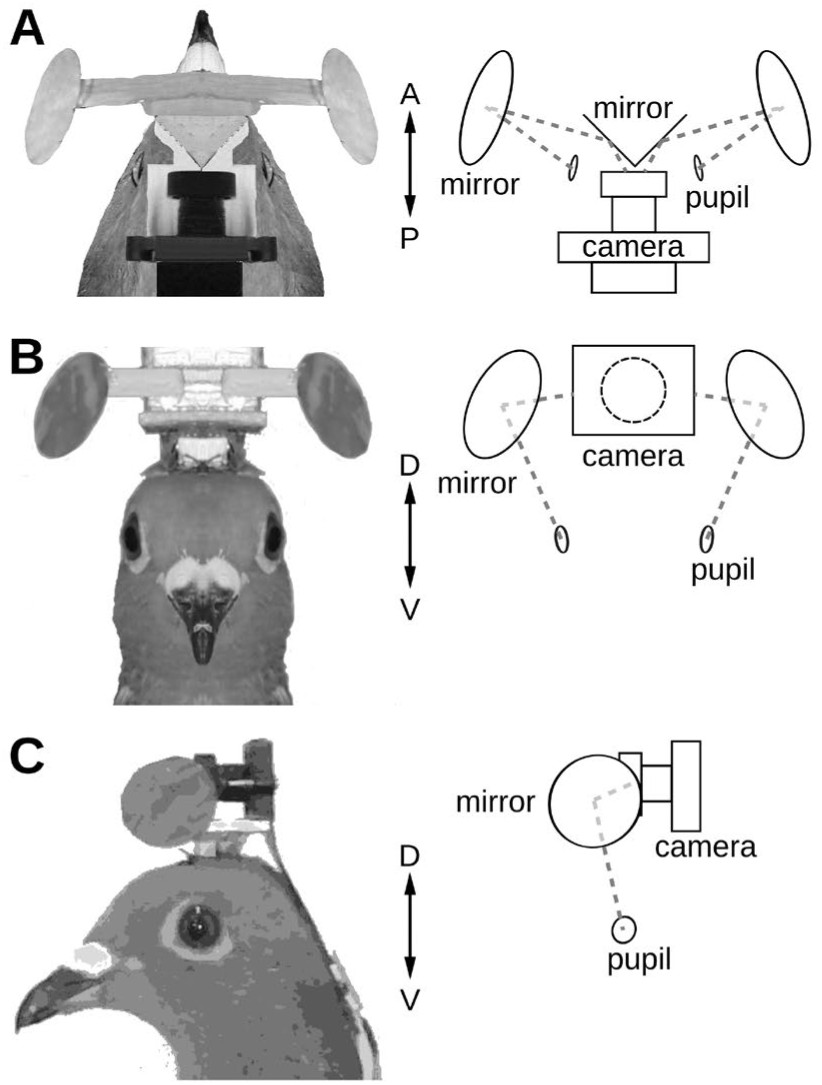
Dual-eye tracking system with head-mounted camera and mirrors. Dorsal (A), frontal (B), and left lateral (C) views of the optical system: Matched photographs (left) and schematics (right). A single camera simultaneously captures both pupils via sets of mirrors.

The saltatory pattern of gaze changes is a classic example of active sensing, in which intentional sensor movements improve perception (Wachowiak, 2011; Skyberg and Niell, 2024). Many walking birds bob their heads, alternating a stable hold phase with a fast, forward thrust phase (Dunlap and Mowrer, 1930; Davies and Green, 1988; Frost, 1978; Wohlschlager *et al*., 1993; Land, 1999). This active sensing strategy helps distinguish external movement and improves estimates of depth perception (Frost, 1978; Troje and Frost, 2000). During non-flight, the fixate-and-saccade gaze strategy is functionally identical whether achieved by eye movements or head rotations. However, during forward flight, two conflicting gaze strategies emerge. Birds can either fixate their eyes within their head or fixate their eyes on their surroundings. By fixating their eyes within their head, birds would perceive the surroundings to move backwards across their retinal images. Or, birds can maintain a stable retinal image on the surroundings by moving their eyes backwards, alternated with fast, corrective forward eye movements (Gioanni, 1988; Kano *et al*., 2018).

Past work has shown that birds stabilize their heads against rotation during flight, and implications for flight control have been well-studied (Bilo *et al*., 1985; Davies and Green, 1988; Warrick *et al*., 2002; Eckmeier *et al*., 2008; Kress *et al*., 2015; Pete *et al*., 2015). Vertebrates have four reflexes that integrate visual and vestibular information to stabilize their head and eyes. Optocollic and optokinetic reflexes stabilize the head and eyes, respectively, by countering optic flow, or movement of the visual scene across the retina. (Gibson, 1958; Koenderink, 1986). Vestibulocollic and vestibulo-ocular reflexes use the semicircular canals and otolith organs to detect head rotations and translations, respectively (Wilson and Melvill Jones, 1979). Birds process visual motion through two parallel neural pathways: the tectofugal system is tuned to small-field object motion, and the accessory optic system (AOS) responds to self-induced translational and rotational optic flow (Simpson, 1984; Iwaniuk and Wylie, 2007). Notably, optic flow neurons have preferred directions in the head frame and share a common frame of reference with the semicircular canals of the vestibular system (Wylie *et al*., 1998). Given the underlying reflexes and visual neural pathways that stabilize head and eye motion under non-flight conditions, we can reasonably expect that birds similarly stabilize their eyes during flight.

Pigeons actively control eye position within the head, which likely allowed for retinal specializations related to head pose (Wallman and Pettigrew, 1985; Nalbach *et al*., 1990; Wallman and Letelier, 1993). Pigeons have two retinal areas of high visual acuity: a central fovea and a dorsally located area dorsalis (Nalbach *et al*., 1990). The central fovea is a lateral-viewing acuity spot close to the optical axis and suited for distant viewing. The dorsally located area dorsalis provides a ventral and frontal acuity spot for nearby viewing. Because of these features, and because head rotation drives most of gaze changes in birds, gaze has been inferred from head orientation during flight (Davies and Green, 1988; Kjaersgaard *et al*., 2008; Eckmeier *et al*., 2008; Kress *et al*., 2015, Ros and Biewener, 2017, Kano *et al*., 2018; Minano *et al*., 2023; Lapsansky *et al*., 2025). However, the assumption that eye movements in flying birds do not change gaze independently from head rotations has not been explicitly tested.

We examined whether pigeons preserve a fixate-and-saccade strategy and stabilize their gaze against their surroundings or maintain their eyes in a head-fixed position as they fly past their surroundings. Using our head-mounted camera-based system, we tracked both eyes of nine pigeons with subsets flying in three different visual environments. We test the hypothesis that flying pigeons accomplish most of their gaze changes via head movements. By simultaneously measuring 3-D head orientation and eye position we determine if gaze can be accurately inferred from head orientation during flight.

## Results

To track bilateral eye positions of free-flying pigeons we constructed a head-mounted optical system with a single camera that captured the pupils of both eyes via two sets of mirrors (Fig. 1; see Methods). We measured eye movements and head rotations in (n=3) pigeons trained to fly to a coop ∼ 100 m away in an outdoor, naturalistic visual environment (Fig. 2A). A light-weight inertial measurement unit recorded head orientation along with the head-mounted optical system that tracked left and right eye positions (Fig. S1). We compared equal 6 s periods of flight and non-flight (Fig. 2A-C). Head rotations and synchronized eye movements alternated periods of stability with rapid saccades (Fig. 2 B, C). In flight, head rotations were 10X larger than eye movements (11.31 ± 1.56 ° vs 1.13 ± 0.19 °, respectively; Fig. 2D, *p* = 0.01, paired-samples *t*-test; n=3). Even though synchronized in time, left and right eye movements did not correlate in magnitude (Fig. 2E; Pearson statistic = 0.14, *p* = 0.24; n=3).

**Fig. 2.**
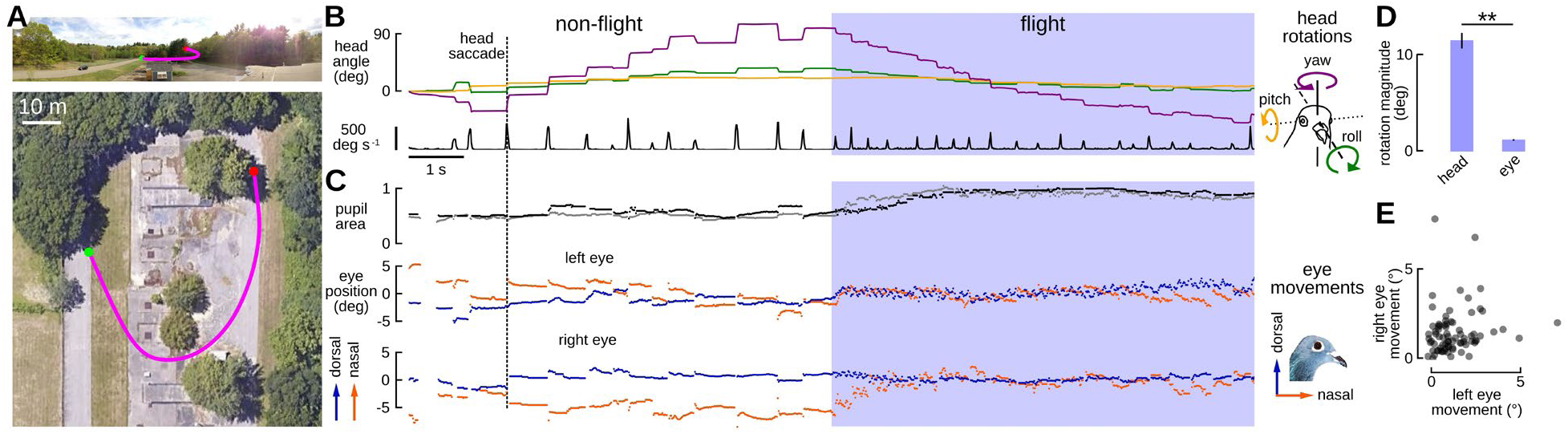
Eye movements are smaller than head rotations during flight. (A) Flight area with a naturalistic visual environment. Top: Horizontal view. Bottom: Vertical view (adapted from Google Maps). Pink trace: Approximate flight trajectory, starting from the green marker. Scale bar: 10 m. (B) Cumulative head rotations throughout an example 6 s of non-flight and 6 s of flight (blue shading). Green: Roll, Orange: Pitch, Purple: Yaw. Bottom black trace represents angular speed of the head, with transients indicating rapid rotations, or head saccades (vertical dotted line illustrates one saccade event). Right inset: Cartoon of the roll, pitch, and yaw axes in the head frame. (C) Time-synchronized recordings of left (grey) and right (black) pupil area (top plot), together with left (middle plot) and right (bottom plot) eye positions plotted along dorsoventral and nasotemporal directions (blue and orange dots, respectively). Right inset: dorsal and nasal directions of eye movements within the head frame. (D) Mean ± SD of individual mean head rotations are larger than eye movements (*p* = 0.01, paired samples *t*-test; n = 3 individuals). (E) The magnitude of saccadic eye movements did not correlate between left and right sides (Pearson correlation coefficient = 0.14, *p* = 0.24; n = 3 individuals).

To characterize eye positioning strategies during flight compared to non-flight, we measured pupil size and eye position for both eyes in nine individual birds distributed across sparse, dense, and naturalistically cluttered visual environments. In all three visual environments, pupil size and eye position followed consistent patterns (Fig. 3A-C, E-G). As pigeons took flight, pupil size increased and eye position relative to the head became less variable (Fig. 3). Pupil area increased in size by 68.9 % during flight compared to non-flight (Fig. 3D, paired samples *t*-test, statistic = -9.65, *p*<0.0001, n=9). When normalized to the 98th percentile pupil size for each pigeon, pupil area during flight averaged 0.88 ± 0.04 versus 0.52 ± 0.09 during non-flight. We compared eye position across individuals by mapping eye position in a head reference frame based on anatomical landmarks. During flight, eye position averaged 3.03 ± 1.26 ° nasal, towards the bill, and 3.39 ± 1.44 ° ventral of the center of the head frame (mean ± SD of individual means). During non-flight, eye position averaged 1.43 ± 2.33 ° nasal, towards the bill, and 4.07 ± 2.37 ° ventral of the center of the head frame (Fig. 3H). Eye position varied less during flight than during non-flight, both along dorsoventral and nasotemporal axes within the head frame (paired samples *t*-test and Wilcoxon signed rank test with Bonferroni-adjusted *p*-values, respectively, with *p* = 0.01 and *p’* = 0.01; Fig. 3H, I, n=9). Within-individual variances in eye position during flight were 0.89 ± 0.17 ° and 0.56 ± 0.19 ° along the nasotemporal and dorsoventral axes, respectively; less than the respective variances in eye position during non-flight at 2.91 ± 1.06 ° and 1.67 ± 1.10 °.

**Fig. 3.**
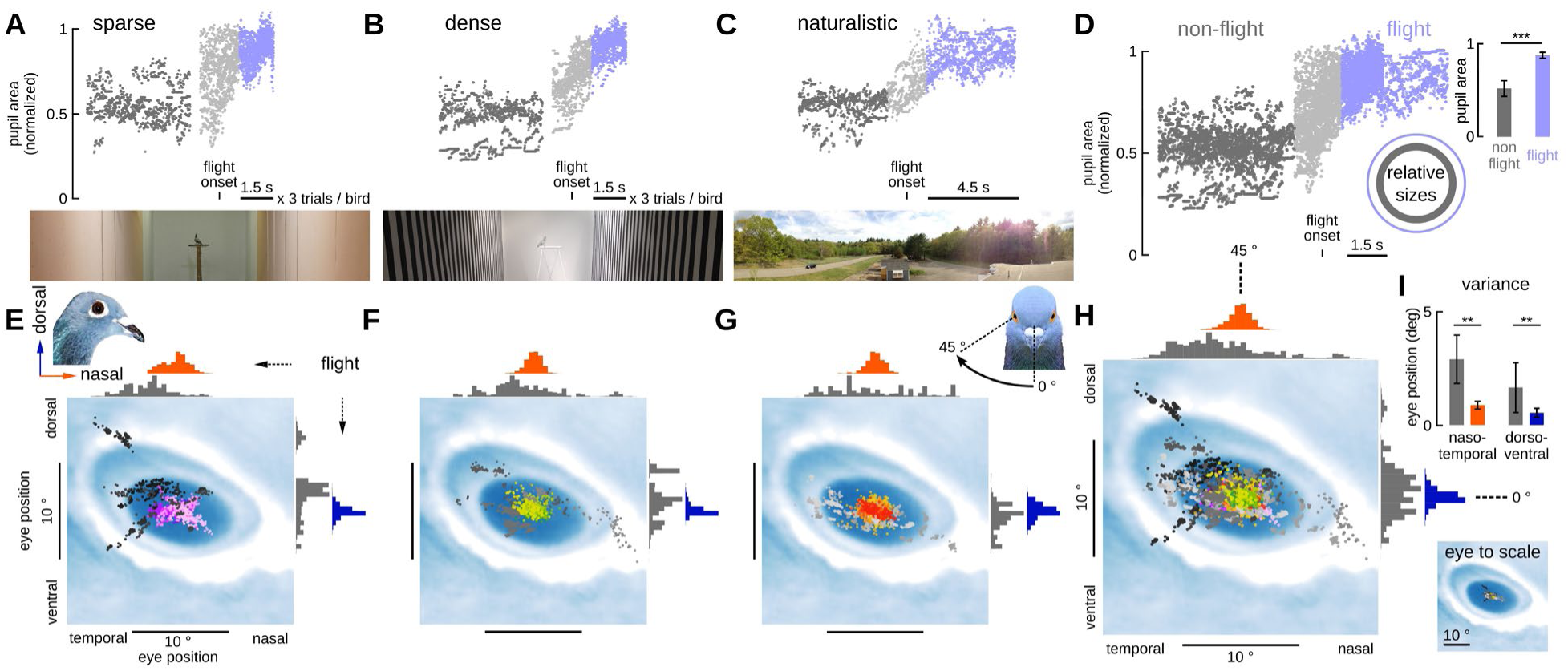
During flight, pupils are enlarged and eye positions are less variable. Pupil areas for left and right eyes during non-flight (4.5 s, dark grey), flight onset (1 s, light grey), and flight (4.5 s, blue) in (A) sparse, (B) dense, and (C) naturalistic visual environments (n = 3 individuals per environment). (A-B) Flight consisted of three trials per individual. (D) Pooled pupil area data across all visual conditions. Bar graph and inset with graphical representation: individual mean pupil area during non-flight (dark grey) and flight (blue; *p* < 0.001, paired *t*-test; mean ± SD). (E) Eye position in the head frame (inset) during non-flight (monochrome shades) and flight (color hues) under (E) cluttered, (F) sparse, and (G) dense visual conditions. Panel background with eye and eye ring orientation for reference (not to scale). Histograms adjacent to eye position plots represent pooled distributions along nasotemporal (top, grey/red) and dorsoventral (right, grey/blue) axes. (G) Inset: graphical representation of the center of the head frame. (H) Pooled eye positions across all individuals and visual environments with associated nasotemporal and dorsoventral histograms. (I) Top: Differences in eye angular position variance between non-flight and flight were assessed using a paired samples *t*-test along the dorsoventral axis and a Wilcoxon signed rank test along the mediolateral axis, with Bonferroni-adjusted *p*-values, *p*’ (*p*’=0.01, n = 9 individuals). Bottom: Eye positions scaled to approximate eye and eye ring orientation.

During flight under both sparse and dense visual conditions, each pigeon moved its eyes to a consistent position relative to their head across trials (Fig. 4). We analyzed eye position in the two indoor visual conditions for three flight trials per individual (Fig. 4A, B; n=6). Individual eye position during flight averaged 3.17 ± 0.37 ° more nasal compared to non-flight, and 1.20 ± 0.09 ° more ventral (Fig. 4A, B). For each individual (n=6), we compared the distance between mean gaze positions of each flight trial to the variance in gaze position during those trials. The individual mean distance between mean flight gaze position across trials (0.92 ± 0.32 °) did not differ from the individual mean variance in flight gaze position within trials (0.79 ± 0.18 °; *t* = 0.79, two-sided *p*-value = 0.44; Fig. 4C). Consequently, across flights and the visual environments examined here, pigeons position their eyes within 1 ° in their head frame, supporting a fixed gaze position during flight.

**Fig. 4.**
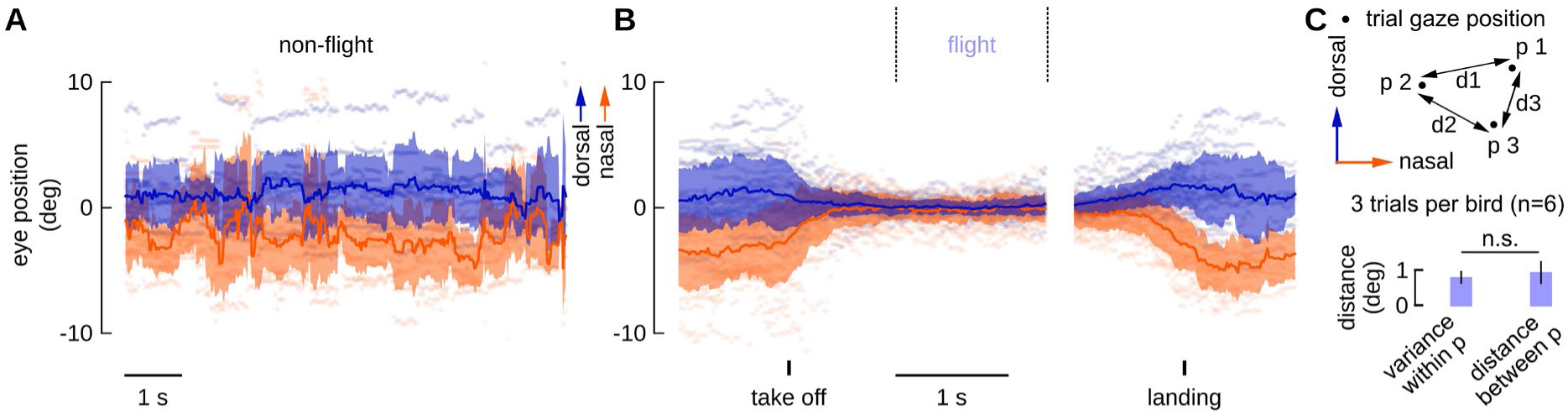
After take-off, eyes move to a flight gaze position. (A) Eye positions of perched pigeons along dorsoventral (blue) and nasotemporal (orange) axes (mean ± SD shading, n = 6 individuals). (B) Eye positions during three flights for each individual, aligned at take-off and landing. (C) Top: Schematic of mean gaze positions during flight for three trials (p1-p3) for an individual, each starting 1 s post-takeoff, with inter-trial Euclidean distances (d1-d3) between those mean gaze positions. Bottom: Comparison of within-trial variance of gaze position and between-trial mean of gaze position distances (*p* = 0.44, two-sample *t*-test, *t* = 0.79, n = 6 individuals); distances between trial gaze positions are indistinguishable from their variance within trials.

## Discussion

Pigeons increase pupil size and fixate their eye position in their head during flight, with reduced eye movement compared to non-flight (Figs. 2, 3). Increased pupil size is consistent with higher temporal acuity demands of vision during flight (Boström *et al*., 2016). In different visual conditions, pigeons consistently position their eyes near the primary horizontal axes of their vestibular systems and preferred directions of optic flow neurons (Figs. 3, 4; Wylie *et al*., 1998). We speculate that this stereotypical eye positioning during flight may help pigeons detect self-motion critical for flight control and navigation (Warren and Wertheim, 2014).

Using a head-mounted system (Fig. 1), we show that the eyes of flying and non-flying pigeons move in synchrony with their rapid head rotations (Fig. 2). In both flight and non-flight conditions, these synchronized eye and head saccades alternate with periods of stable eye position and head orientation (Fig. 2). However, in-flight eye movements are an order of magnitude smaller than head rotations (Fig. 2). In prior research on walking or restrained birds, the head and eyes also rotate in synchrony, where eye movements augment stabilization of the visual scene on the retina (Pratt, 1982; Wallman and Letelier, 1993; Wohlschlager *et al*., 1993; Maurice and Gioanni, 2004; Lapsansky *et al*., 2025). Comparatively, flies similarly stabilize vision by rotating their head in synchrony with body movements, and further reduce optic flow with fine eye movements (Land, 1973; Schilstra and Van Hateren, 1998, Hengstenberg *et al*. 1986; Nalbach and Hengstenberg, 1994; Cellini *et al*., 2021, Fenk *et al*., 2022).

When flying across sparse, dense, and naturalistically cluttered visual environments, pigeons increased their pupil size and displayed smaller, less variable eye movements within a narrower range of eye positions (Figs. 2, 4). Increased pupil size indicates increased sympathetic activity (Bradshaw, 1967; Bradley et al., 2008). Functionally, a larger pupil lets in more light, allowing for shorter photoreceptor response times, consistent with higher information processing demands during flight (Dodt and Wirth, 1953; Hendricks, 1966). Varying optic flow conditions in different environments can elicit differing responses in birds depending on behavioral context (Maurice *et al*., 2006; McArthur and Dickman, 2011), proximity of surroundings (Bhagavatula *et al*., 2011; Dakin *et al*., 2016), or composition of the sensory environment, such as demonstrated in flies (Rimniceanu *et al*., 2024). We occasionally observed nystagmic eye movements consistent with the optokinetic reflex where both eyes move backwards, to track the wide-field surround, and rapidly reset nasally (Fig. 2 C, last 2 s of flight; Fig. 4B; Gioanni, 1988; Maurice and Gioanni, 2004). However, as flight speed increased during flight trials, optic flow speed likely increased. Given higher optic flow speeds, these optokinetic eye movements diminished, and eye positions converged to a consistent location relative to the head (Figs. 2-4). For these flights, during which the birds flew solo and without obstacles, a consistent fixed eye positioning indicates a general visual strategy during flight at relatively high optic flow speeds (Fig. 3E-H). In flying pigeons, head rotations seem to drive most of gaze changes, with the eyes nearly locked within the head (Figs. 2-4).

Pigeons also substantially constrain the range and variability of their eye movements, adopting a stereotyped flight gaze position during flight (Figs. 3, 4). By representing eye positions in a head-centered reference frame based on anatomical landmarks, we can compare eye position across individuals. Pigeons consistently return their eyes to the same in-flight position across flight trials. Following take-off, the eyes rotate slightly forwards along a nasotemporal angle of ∼45 ° to the head midline, before returning more laterally upon landing (Fig. 4B).

The observed eye positioning aligns with the primary horizontal axis of neural representations in both visual and vestibular sensory systems (Wylie and Frost, 1999; Wylie, 2013). In vertebrates, vestibular afferents project to the cerebellum and brain stem nuclei (Warren and Wertheim, 1990), which relay signals to oculomotor nuclei. Neurons encoding translational and rotational optic flow have preferred directions in the head reference frame along three axes: one vertical and two horizontal axes oriented ±45 ° from the midline. Both visual and vestibular systems are maximally sensitive at ±45 ° along the nasotemporal azimuth, indicating that optic flow neurons share a common reference frame with the semicircular canals (Wylie *et al*., 1998).

Aligning the lateral fovea with the primary horizontal axis of multimodal sensory representation offers several non-mutually exclusive functional advantages during flight. Based on the common reference frame between neural representations of vision and mechanosensation, retinotopic optic flow fields could be transformed into head-frame coordinates, or eyes could be actively kept in line with the primary horizontal axis of the vestibular system (Zipser and Anderson, 1988). Physically aligning the sensory systems by adopting a flight gaze position within the head may facilitate representing visual and vestibular information in a single reference frame.

Aligning reference frames may also facilitate sensor fusion between vision and mechanosensation. Sensor fusion can reduce measurement uncertainty in estimates of self-motion (Angelaki *et al*., 2011). The vestibular system could register the magnitude of rapid head saccades to reduce the potential error that would originate from the visual system, particularly at higher rotation speeds (DeAngelis and Anglaki, 2012). Similar advantages of sensor fusion have been observed in diptera, where vision and mechanosensation complement and overlap detection of self-motion across the frequency spectrum (Hengstenberg, 1991, Sherman and Dickinson, 2002; Huston and Krapp, 2009).

Stabilizing the eye within the head while simultaneously stabilizing the head in space reduces the ambiguity of retinal image motion. When the eye moves less than a so-called ‘Just Noticeable Difference’ threshold (JND; MacKay, 1973), all optic flow can be perceived as self-motion, rather than eye motion within the orbit (Warren and Wertheim, 1990). Effectively, stabilizing gaze against rotations isolates the optic flow most relevant to spatial navigation.

It remains an open question whether the observed alignment represents an active sensing strategy that serves to speed up neural computations through simplification or reflects a spurious correlation that arises from independent optimization of each sensory system. Explicit experimental manipulation of the sensory alignment during flight could provide further clarity in the functional roles of the putative flight gaze position.

## Materials and Methods

### Subjects

We housed homing and wild-type pigeons (*Columba livia*; age 1-2 years; n = 9 total; mean ± SD body mass: 0.45 ± 0.05 kg) at the Concord Field Station (Bedford, MA). Three individuals from a homing group were trained to return to their coop upon outdoor release. The remaining six individuals were trained to fly in the indoor setups. This study was carried out in accordance with the guidelines of Harvard University’s Institutional Animal Care and Use Committee.

### Subject training

For indoor experiments, male and female pigeons were individually trained to fly between two perches placed on either end of a 22-m-long by 2-m-wide by 2.4-m-high flight corridor. Each individual was induced to take off from a perch by the experimenter approaching. A training procedure was considered complete when an individual flew consistently between perches upon approach-command. Training required approximately 100 flights, distributed over 5-8 days. For outdoor experiments, male homing pigeons from a male-female pair were taken out of their homing coop, hand-released from a distance of approximately 25 m and allowed to return. The procedure was repeated for 2 - 3 weeks until the pigeons were trained to consistently return to their coop. For outdoor experiments, three individuals were released approximately 100 m from the homing coop.

### Indoor experimental setup

The pigeons were flown indoors in a custom-built flight corridor in a building temperature-controlled between 23-25° C. The walls of the corridor were painted with white paint (white, low odor; Behr, Santa Ana, USA) and decorated with vertical 0.1 m wide, black, machine cut cardboard stripes (Canford paper, Daler Rowney, UK), separated by 0.1 m to form a square-wave pattern. The floor of the corridor was similarly covered with butcher paper and black stripes, arranged parallel to match those found on the walls. Illumination was provided by 250 and 500 W, flicker-free, halogen lights (HDX, Nashville, TN).

### Camera-based eye-tracking system

The head stage of the eye-tracking system comprised a balsa wood base, on which a triangular balsa wood prism supported circular mirrors facing the camera lens at a 45 ° angle (Fig.1, S1). Two mirrors mounted directly above the pigeon’s eyes with balsa wood doweling and projected the image of the eyes through the mirrors mounted on the triangular prism into the camera (see Fig. 1). We used a modified Mobius Action Camera that acquired images at 60 Hz (Huizhou tuopu xunshi Technology Co., Ltd, PR China) with a Wingcam lens (Hobbyking.com). The mirrors were made from 12 mm diameter glass cover slips (SKU 1943-10012; Bellco Glass, Vineland, NJ), custom-mirrored using a silver kit (Mini Silver Kit, Forest Park, IL, USA). The head stage was mounted using hot-melt adhesive (Gorilla Glue, Cincinnati, OH, USA) on two bass wood posts. The posts were separated by 3 mm along the mid-sagittal plane and securely mounted on the skull of each individual (see next section). The camera controller board (Mobius camera controller) and 400 mAh lithium-ion polymer battery (Data Power Technology Ltd., Shenzhen City, PR China) were attached to a bass wood baseplate, which was secured onto elastic adhesive tape (Elastikon, New Brunswick, NJ, USA) affixed to the dorsal side of the individual at the mantle-back transition. The mass of the head-stage and body-stage totaled 1.25 g and 27 g, respectively (Fig. S2).

### Surgical procedure

Two bass wood posts were mounted onto the skull using UV curable adhesive. To perform surgery, the subject was placed under general anesthesia using Isoflurane gas (2 %). We mounted the 4 mm long, 3.2 mm edge posts vertically on the skull using UV-cure adhesive (Loctite AA 349). A fore-aft incision exposed the skull, positioned mid-sagitally on the dorsal aspect of the frontal bone of the skull, centered along the AP axis at the posterior end of the sclerotic ring. The exposed skull was cleaned using surgical gauze prior to removal of subdermal connective tissue. To improve adhesion, we cleaned and desiccated the exposed bone tissue with 70% isopropyl alcohol (S&S brand, LLC). The incision was closed around the posts using 8-0 Vicryl sutures (polyglactin 910, Ethicon Inc.). To reduce discomfort, we intra-muscularly administered a standard dose of analgesic (Flunixin meglumine, 4mg/kg). Following experiments, we removed the posts and adhesive under similar anesthesia by a twisting moment about the DV axis that easily dislodged the adhesive from the skull. We fully closed the incision with additional sutures and returned subjects to their coop for recuperation.

### Orientation sensor

For outside flight, an orientation sensor tracked head orientation (Fig. 2). We used a 9 degrees-of-freedom Absolute Orientation Sensor BNO055 (Bosch Sensortec GmbH, Germany) mounted on an Adafruit industry break-out board, powered by a 110 mAh lithium-ion polymer battery.

The BNO055 connected to an Arduino pro mini (Qualcomm, San Diego, CA, USA) via an I^2^C bus. The Arduino pro mini was connected to an Open Log data logger (Sparkfun Electronics, USA) using a serial port. The Open Log data logger saved data to a 16 GB micro-SD card (Toshiba, Japan). Orientation sensor data were saved as three nested Euler angles at 100 Hz. Euler angles were used to reconstruct the rotation matrix from global to sensor, and, thus, head frame. Head frame orientation was integrated over time to give cumulative head rotations about roll, pitch, and yaw axes (see Fig. 2).

### GPS tracking of flights

A custom-built GPS tracker (NavSpark, Taiwan) was used to track the outdoor flight of individual pigeons to obtain estimates of flight trajectory statistics, such as path length and flight speed. The GPS tracker sampled at 20 Hz and was powered by a 400mAh Lithium-ion polymer battery. Two GPS-tracked flight trajectories were used to approximate representative flight paths (Fig. 2A).

### Data analysis

Analyses were based on bilateral eye-tracking over a balanced 4.5 s of non-flight and 4.5 s of flight for each individual (n=9). We semi-automatically tracked pupil position and size using image luminance thresholding in custom-written scripts in Matlab (The MathWorks, Inc., Natick, MA). To optimize pupil contrast across varying lighting conditions, red, green, and blue channel values were re-stretched within a region of interest surrounding each eye. Following an automatic fit of an ellipses on binarized images of the eye, manual verification removed erroneous tracking frames, predominantly during blinking. To compare changes in pupil size between non-flight to flight, pupil size was normalized to the 98^th^ percentile across all recordings for each individual.

We expressed tracked eye positions in a head coordinate system based on anatomical landmarks, independent of eye positions, allowing for comparisons of gaze across individuals. We oriented the head-refence frame by defining the temporo-nasal axis as 34 ° upwards (Erichsen, 1989) relative to a line from the temporal to the nasal canthi of the orbital ring. The dorso-ventral axis was defined as normal to the temporo-nasal axis. For each eye of each individual, the head reference frame was normalized to its iris diameter measured separately with calipers during the surgical procedure. For all data included for analysis, every trial for a given individual was inspected to ensure that mirror positioning did not change either within or across trials.

### Quantification and statistical analyses

We tracked eyes with custom scripts (Matlab) and analyzed data using Python (python.org; matplotlib.org). Sample sizes, n, refer to the number of individual birds tested. Variance across individuals is represented by standard deviation of individual means (Figs. 2D, 3D, I, 4A-C). Statistical differences between individual mean head rotations and eye movements and between pupil area during flight and non-flight were assessed with paired samples *t*-tests (Fig. 2D, 3D, respectively). Covariance between left and right saccadic eye movements was assessed using the Pearson Correlation Coefficient (Fig. 2E). Differences in variance of pupil angular position between non-flight and flight were assessed using a paired samples *t*-test along the dorsoventral axis and a Wilcoxon signed rank test along the mediolateral axis with Bonferroni-adjusted *p*-values (Fig. 3I). Differences between within-flight variance of gaze position and between-flight mean gaze position distances were assessed with a two-sample *t*-test (Fig. 4C).

## Acknowledgments

We thank P. A. Ramirez for animal care and are grateful to B. Frost for insightful discussions and guiding our work. We further appreciate helpful contributions from H-T Lin, A. N. Ahn, and A. M. Mountcastle. This study was supported by a grant from the Office of Naval Research (N0014-10-1-0951) to A.A.B.

## Author contributions

Conceptualization, I.G.R. and A.A.B.; methodology, I.G.R., P.S.B, and A.A.B.; software, I.G.R; validation, I.G.R and P.S.B.; formal analysis, I.G.R; formal analysis, I. G. R.; performed research, I.G.R. and P.S.B; resources, I.G.R., P.S.B., and A.A.B.; writing – original draft, I.G.R and P.S.B.; writing – review & editing, I.G.R., P.S.B, and A.A.B.; funding: A.A.B.

## Competing Interests

The authors declare no competing interests.

## Data and materials availability

The eye tracking software is available at http://github.com/flivo/Cl_pupiltrack. All data in main text figures and their associated supplementary figures have been deposited on Mendeley at https://doi.org/10.17632/h3ns2x976d.1 and will be publicly available upon the date of publication. All original code to process the data has also been deposited on Mendeley at https://doi.org/10.17632/h3ns2x976d.1. Any additional information required to reanalyze the data reported in this paper is available from the lead contact upon request.

## Notes

https://doi.org/10.17632/h3ns2x976d.1

### Competing Interest Statement

The authors have declared no competing interest.

